## Supplementary material for "Finding New Molecular Targets of Familiar Natural Products Using In Silico Target Prediction"

### CONTENT

S-1 PHARMACOPHORE MODELS USED IN THIS STUDY: 4

S-2 PIPELINE PILOT PROTOCOL: 17

S-3 FILE SCHEME: 18

S-4 LIGAND PROFILER PARAMETER SETTINGS 19

S-5 QUALITY CONTROL OF 1 – 10: 20

- Phloretin (1) CAS: 60-82-2 20
- 3-OH-phloretin (2) CAS: 57765-66-9 20
- 2',6'-dihydroxy-4'-methoxy DHC (3) CAS: 35241-55-5 21
- Asebogenin (4) CAS: 35241-54-4 21
- Calomelanen (5) CAS: 520-42-3 22
- Sieboldin (6) CAS: 18777-73-6 22
- Phloridzin (7) CAS: 60-81-1 23
- Trilobatin (8) CAS: 4192-90-9 23
- Phloretin-2'-xyloglucoside (9) CAS: 145758-09-4 24
- Neohesperidin DHC (10) CAS: 20702-77-6 24

S-6 POSITIVE CONTROLS: 25

S-7 REFERENCES: 26

### S-1 PHARMACOPHORE MODELS USED IN THIS STUDY:

Table S 1. Pharmacophore Models used throughout this study.

| Model Name | Software | UniProt entry | Ref. |
|---|---|---|---|
| model_DS4 | DS | 3BHS1_HUMAN | (Kurzemann, 2016) |
| Model_1 | LS | 3BHS1_HUMAN | (Kurzemann, 2016) |
| Model_2 | LS | 3BHS1_HUMAN | (Kurzemann, 2016) |
| AChE-1acj-tha-HBA-vers | DS | ACES_HUMAN | (Rollinger et al., 2009) |
| AChE-1acj-tha-xvols | DS | ACES_HUMAN | (Rollinger et al., 2009) |
| AChE-GNT-Hypo1-shape | DS | ACES_HUMAN | (Rollinger et al., 2009) |
| AChE-X015-1 | DS | ACES_HUMAN | (Rollinger et al., 2009) |
| 17b-HSD5-1ry8-rut-1.60-x-1 | DS | AK1C3_HUMAN | (Schuster et al., 2011a) |
| 17b-HSD5-1ry8-rut-1.60-x-1-s | DS | AK1C3_HUMAN | (Schuster et al., 2011a) |
| 17b-HSD5-1s2a-imn-1.70-x-1 | DS | AK1C3_HUMAN | (Schuster et al., 2011a) |
| 17b-HSD5-1s2a-imn-1.70-x-1-s | DS | AK1C3_HUMAN | (Schuster et al., 2011a) |
| 17b-HSD5-1s2c-flf-1.80-x-1 | DS | AK1C3_HUMAN | (Schuster et al., 2011a) |
| 17b-HSD5-1s2c-flf-1.80-x-1-s | DS | AK1C3_HUMAN | (Schuster et al., 2011a) |
| 17b-HSD5-1s2c-flf-1.80-x-2-s | DS | AK1C3_HUMAN | (Schuster et al., 2011a) |
| 17b-HSD5-1zq5-e04-1.30-x-1 | DS | AK1C3_HUMAN | (Schuster et al., 2011a) |
| 17b-HSD5-1zq5-e04-1.30-x-1-s | DS | AK1C3_HUMAN | (Schuster et al., 2011a) |
| 17b-HSD5-1zq5-e04-1.30-x-2-s | DS | AK1C3_HUMAN | (Schuster et al., 2011a) |
| 17b-HSD5-2f38-15m-2.00-x-1-s | DS | AK1C3_HUMAN | (Schuster et al., 2011a) |
| 17b-HSD5-X009-1 | DS | AK1C3_HUMAN | (Schuster et al., 2011a) |
| 17b-HSD5-X009-2 | DS | AK1C3_HUMAN | (Schuster et al., 2011a) |
| FLAPophore6exclu_LS3-03 | LS | AL5AP_HUMAN | (Temml et al., 2017) |
| FLAPophore7exclu_LS3-03 | LS | AL5AP_HUMAN | (Temml et al., 2017) |
| LS3.12-CYP11B1_2T_D-1-8 | LS | C11B1_HUMAN | (Akram et al., 2019) |

| | | | |
|---|---|---|---|
| LS3.12-CYP11B1_2T_D-1-8_refined-Veronika | LS | C11B1_HUMAN | (Akram et al., 2019) |
| LS3.12-CYP11B2_C-1-5 | LS | C11B2_HUMAN | (Akram et al., 2019) |
| LS3.12-CYP11B2_C-1-5-refined-Veronika | LS | C11B2_HUMAN | (Akram et al., 2019) |
| CYP17_3RUK_LS4_03_MA | LS | CP17A_HUMAN | * |
| CYP17_Abi_Orteronel_LS4_03_MA | LS | CP17A_HUMAN | * |
| aromatase-X006-1 | DS | CP19A_HUMAN | (Schuster et al., 2006) |
| aromatase-X006-2 | DS | CP19A_HUMAN | (Schuster et al., 2006) |
| aromatase-X006-3 | DS | CP19A_HUMAN | (Schuster et al., 2006) |
| CYP2D6-quant-X012-1 | DS | CP2D6_HUMAN | (Hochleitner et al., 2017) |
| CYP2D6-X012-2 | DS | CP2D6_HUMAN | (Hochleitner et al., 2017) |
| CYP2D6-X012-3 | DS | CP2D6_HUMAN | (Hochleitner et al., 2017) |
| CYP2D6-X012-41 | DS | CP2D6_HUMAN | (Hochleitner et al., 2017) |
| CYP2D6-X012-42 | DS | CP2D6_HUMAN | (Hochleitner et al., 2017) |
| LS3-03b-17b-HSD2-model6-refined | LS | DHB2_HUMAN | (Vuorinen et al., 2017) |
| LS3-03b-17b-HSD2-specific-model-12 | LS | DHB2_HUMAN | (Vuorinen et al., 2017) |
| LS3-03b-17b-HSD2-specific-model-6 | LS | DHB2_HUMAN | (Vuorinen et al., 2017) |
| LS3-03b-17b-HSD2-specific-model-8 | LS | DHB2_HUMAN | (Vuorinen et al., 2017) |
| 17b-HSD3-DS-model-1-X007-1 | DS | DHB3_HUMAN | (Schuster et al., 2011a) |
| 17b-HSD3-DS-model-2-X007-31 | DS | DHB3_HUMAN | (Schuster et al., 2011a) |
| 17b-HSD3-LS3-1-317364-65-1-and-317364-71-9-model-4 | LS | DHB3_HUMAN | (Schuster et al., 2011a) |
| 17b-HSD3-LS3-1-317364-65-1-and-317364-71-9-model-5 | LS | DHB3_HUMAN | (Schuster et al., 2011a) |
| 17b-HSD3-LS3-1-875895-55-9-875895-40-2-model-1 | LS | DHB3_HUMAN | (Schuster et al., 2011a) |
| 17b-HSD3-LS3-1-Benzophenone-1-919118-27-7-model-1 | LS | DHB3_HUMAN | (Schuster et al., 2011a) |
| 17b-HSD3-LS3-1-Harada-2012-19-Fink-18m-model-4 | LS | DHB3_HUMAN | (Schuster et al., 2011a) |
| 17b-HSD3-LS3-1-Harada-2012-19-Fink-18m-model-7 | LS | DHB3_HUMAN | (Schuster et al., 2011a) |
| 17b-HSD4-quercetin-Wetzel-final | DS | DHB4_HUMAN | (Schuster et al., 2011a) |

| | | | |
|---|---|---|---|
| 17b-HSD4-Quercetin-Wetzel-HBA-F-model3-2 | DS | DHB4_HUMAN | (Schuster et al., 2011a) |
| 17b-HSD4-Quercetin-Wetzel-HBA-F-model8-1 | DS | DHB4_HUMAN | (Schuster et al., 2011a) |
| 17b-HSD4-Quercetin-Wetzel-model-4-final-LS3-1 | LS | DHB4_HUMAN | (Schuster et al., 2011a) |
| 17b-HSD7-Fiala-DS-shapemodell_30 | DS | DHB7_HUMAN | (Fiala, 2017) |
| 17b-HSD7-Fiala-DS-shapemodell_42 | DS | DHB7_HUMAN | (Fiala, 2017) |
| 17b-HSD7-Fiala-DS-shapemodell_43 | DS | DHB7_HUMAN | (Fiala, 2017) |
| 17b-HSD7-Fiala-DS-shapemodell_49 | DS | DHB7_HUMAN | (Fiala, 2017) |
| LS4-09-17b-HSD7-Fiala-model2-1 | LS | DHB7_HUMAN | (Fiala, 2017) |
| LS4-09-17b-HSD7-Fiala-model4-1 | LS | DHB7_HUMAN | (Fiala, 2017) |
| 11b-HSD-1-refinedX005-1-HBA-model4new | DS | DHI1_HUMAN | (Vuorinen et al., 2013) |
| 11b-HSD1-X005-1-HBA-ohneShape-model4 | DS | DHI1_HUMAN | (Vuorinen et al., 2013) |
| 11b-HSD1-X005-1-model1 | DS | DHI1_HUMAN | (Vuorinen et al., 2013) |
| 11b-HSD-triterpenes-common-features | DS | DHI1_HUMAN | (Vuorinen et al., 2013) |
| 11b-HSD2-refinedX005-2-model2new | DS | DHI2_HUMAN | (Denise V. Kratschmar et al., 2011a) |
| 11b-HSD2-X005-2-model2 | DS | DHI2_HUMAN | (Denise V. Kratschmar et al., 2011a) |
| LS3-03a-11bHSD2-model3 | LS | DHI2_HUMAN | (Denise V. Kratschmar et al., 2011a) |
| LS3-03a11bHSD2-model5 | LS | DHI2_HUMAN | (Denise V. Kratschmar et al., 2011a) |
| LS3-03a-refined-11b-HSD2model-model3new | LS | DHI2_HUMAN | (Denise V. Kratschmar et al., 2011a) |
| LS3-03a-refined-11bHSD2-model-model5new | LS | DHI2_HUMAN | (Denise V. Kratschmar et al., 2011a) |
| ERa-antagonist-LS3-01-model584 | LS | ESR1_HUMAN | (Brunner, 2011) |
| ERa-antagonist-LS3-01-model585 | LS | ESR1_HUMAN | (Brunner, 2011) |
| ERa-antagonist-LS3-01-model586 | LS | ESR1_HUMAN | (Brunner, 2011) |
| ER-agonist-LS3-01-model009 | LS | ESR1_HUMAN | (Brunner, 2011) |
| ER-agonist-LS3-01-model022 | LS | ESR1_HUMAN | (Brunner, 2011) |
| ER-agonist-LS3-01-model030 | LS | ESR1_HUMAN | (Brunner, 2011) |
| ER-agonist-LS3-01-model040 | LS | ESR1_HUMAN | (Brunner, 2011) |
| ER-agonist-LS3-01-model042 | LS | ESR1_HUMAN | (Brunner, 2011) |
| ER-agonist-LS3-01-model047 | LS | ESR1_HUMAN | (Brunner, 2011) |
| ER-agonist-LS3-01-model052 | LS | ESR1_HUMAN | (Brunner, 2011) |
| ER-agonist-LS3-01-model062 | LS | ESR1_HUMAN | (Brunner, 2011) |
| ER-agonist-LS3-01-model069 | LS | ESR1_HUMAN | (Brunner, 2011) |

| | | | |
|---|---|---|---|
| ER-agonist-LS3-01-model073 | LS | ESR1_HUMAN | (Brunner, 2011) |
| ER-agonist-LS3-01-model084 | LS | ESR1_HUMAN | (Brunner, 2011) |
| ER-agonist-LS3-01-model087 | LS | ESR1_HUMAN | (Brunner, 2011) |
| ER-agonist-LS3-01-model099 | LS | ESR1_HUMAN | (Brunner, 2011) |
| ER-antagonist-LS3-01-model567 | LS | ESR1_HUMAN | (Brunner, 2011) |
| ER-antagonist-LS3-01-model571 | LS | ESR1_HUMAN | (Brunner, 2011) |
| ER-antagonist-LS3-01-model572 | LS | ESR1_HUMAN | (Brunner, 2011) |
| ER-antagonist-LS3-01-model573 | LS | ESR1_HUMAN | (Brunner, 2011) |
| ER-antagonist-LS3-01-model574 | LS | ESR1_HUMAN | (Brunner, 2011) |
| ER-antagonist-LS3-01-model575 | LS | ESR1_HUMAN | (Brunner, 2011) |
| ER-antagonist-LS3-01-model577 | LS | ESR1_HUMAN | (Brunner, 2011) |
| ER-antagonist-LS3-01-model578 | LS | ESR1_HUMAN | (Brunner, 2011) |
| ER-antagonist-LS3-01-model580 | LS | ESR1_HUMAN | (Brunner, 2011) |
| ER-antagonist-LS3-01-model583 | LS | ESR1_HUMAN | (Brunner, 2011) |
| ER-antagonist-LS3-01-model587 | LS | ESR1_HUMAN | (Brunner, 2011) |
| ERb-agonist-LS3-01-model100 | LS | ESR2_HUMAN | (Brunner, 2011) |
| ERb-agonist-LS3-01-model102 | LS | ESR2_HUMAN | (Brunner, 2011) |
| ERb-agonist-LS3-01-model103 | LS | ESR2_HUMAN | (Brunner, 2011) |
| ERb-agonist-LS3-01-model104 | LS | ESR2_HUMAN | (Brunner, 2011) |
| ERb-antagonist-LS3-01-model567 | LS | ESR2_HUMAN | (Brunner, 2011) |
| ERb-antagonist-LS3-01-model571 | LS | ESR2_HUMAN | (Brunner, 2011) |
| ERb-antagonist-LS3-01-model572 | LS | ESR2_HUMAN | (Brunner, 2011) |
| ERb-antagonist-LS3-01-model573 | LS | ESR2_HUMAN | (Brunner, 2011) |
| ERb-antagonist-LS3-01-model574 | LS | ESR2_HUMAN | (Brunner, 2011) |
| ERb-antagonist-LS3-01-model575 | LS | ESR2_HUMAN | (Brunner, 2011) |
| ERb-antagonist-LS3-01-model577 | LS | ESR2_HUMAN | (Brunner, 2011) |
| ERb-antagonist-LS3-01-model578 | LS | ESR2_HUMAN | (Brunner, 2011) |
| ERb-antagonist-LS3-01-model580 | LS | ESR2_HUMAN | (Brunner, 2011) |
| ERb-antagonist-LS3-01-model583 | LS | ESR2_HUMAN | (Brunner, 2011) |
| ERb-antagonist-LS3-01-model587 | LS | ESR2_HUMAN | (Brunner, 2011) |
| GR-1m2z-dex-2.50-T-1 | DS | GCR_HUMAN | (Linder, 2010) |

| | | | |
|---|---|---|---|
| GR-1nhz-486-2.30-T-1 | DS | GCR_HUMAN | (Linder, 2010) |
| GR-1nhz-486-2.30-T-2 | DS | GCR_HUMAN | (Linder, 2010) |
| GR-1nhz-486-2.30-T-3 | DS | GCR_HUMAN | (Linder, 2010) |
| GR-1nhz-486-2.30-T-4 | DS | GCR_HUMAN | (Linder, 2010) |
| GR-1nhz-486-2.30-T-5 | DS | GCR_HUMAN | (Linder, 2010) |
| GR-1p93-dex-2.70-T-1 | DS | GCR_HUMAN | (Linder, 2010) |
| GR-3bqd-day-2.50-T-1 | DS | GCR_HUMAN | (Linder, 2010) |
| GR-3bqd-day-2.50-T-2 | DS | GCR_HUMAN | (Linder, 2010) |
| GR-3cld-gw6-2.84-T-1 | DS | GCR_HUMAN | (Linder, 2010) |
| GR-3e7c-866-2.15-T-1 | DS | GCR_HUMAN | (Linder, 2010) |
| GR-T001-1 | DS | GCR_HUMAN | (Linder, 2010) |
| GR-T001-2 | DS | GCR_HUMAN | (Linder, 2010) |
| GR-T001-3 | DS | GCR_HUMAN | (Linder, 2010) |
| GR-T002-1 | DS | GCR_HUMAN | (Linder, 2010) |
| GR-T002-2 | DS | GCR_HUMAN | (Linder, 2010) |
| GR-T003-1 | DS | GCR_HUMAN | (Linder, 2010) |
| GR-T004-1 | DS | GCR_HUMAN | (Linder, 2010) |
| GR-T005-1 | DS | GCR_HUMAN | (Linder, 2010) |
| GR-T006-1 | DS | GCR_HUMAN | (Linder, 2010) |
| GR-T007-1 | DS | GCR_HUMAN | (Linder, 2010) |
| GR-T008-1 | DS | GCR_HUMAN | (Linder, 2010) |
| GR-T009-1 | DS | GCR_HUMAN | (Linder, 2010) |
| GR-T010-1 | DS | GCR_HUMAN | (Linder, 2010) |
| GR-T011-1 | DS | GCR_HUMAN | (Linder, 2010) |
| GR-T012-1 | DS | GCR_HUMAN | (Linder, 2010) |
| GR-T013-1 | DS | GCR_HUMAN | (Linder, 2010) |
| GR-T014-1 | DS | GCR_HUMAN | (Linder, 2010) |
| GR-T015-1 | DS | GCR_HUMAN | (Linder, 2010) |
| GR-T015-2 | DS | GCR_HUMAN | (Linder, 2010) |
| GR-T016-1 | DS | GCR_HUMAN | (Linder, 2010) |
| GR-T017-1 | DS | GCR_HUMAN | (Linder, 2010) |

| | | | |
|---|---|---|---|
| sEH-C12_1_DS3.5 | DS | HYES_HUMAN | (Waltenberger et al., 2016) |
| sEH-C15_1DS3.5 | DS | HYES_HUMAN | (Waltenberger et al., 2016) |
| sEH-1VJ5mod+3ANTmod_merged(Ref.3ANT)_LS_3-02 | LS | HYES_HUMAN | (Waltenberger et al., 2016) |
| sEH-1ZD5_mod5_LS_3-02 | LS | HYES_HUMAN | (Waltenberger et al., 2016) |
| sEH-3ANT[A]_modifiziert_LS_3-02 | LS | HYES_HUMAN | (Waltenberger et al., 2016) |
| sEH-3ANT+3ANTmod_merged_LS_3-02 | LS | HYES_HUMAN | (Waltenberger et al., 2016) |
| sEH-3I1Y_mod2_LS_3-02 | LS | HYES_HUMAN | (Waltenberger et al., 2016) |
| sEH-3I1Y_mod4_LS_3-02 | LS | HYES_HUMAN | (Waltenberger et al., 2016) |
| sEH-3KOO_mod2_LS_3-02 | LS | HYES_HUMAN | (Waltenberger et al., 2016) |
| sEH-3KOO_mod5_LS_3-02 | LS | HYES_HUMAN | (Waltenberger et al., 2016) |
| sEH-3OTQ_mod4_LS_3-02 | LS | HYES_HUMAN | (Waltenberger et al., 2016) |
| sEH-Cl2_Shared_mod1_LS_3-02 | LS | HYES_HUMAN | (Waltenberger et al., 2016) |
| 5-LO-N004-1 | DS | LOX5_HUMAN | (Noha, 2013) |
| 5-LO-N004-2 | DS | LOX5_HUMAN | (Noha, 2013) |
| 5-LO-W001-5 | DS | LOX5_HUMAN | (Walzl, 2010) |
| LS4-09-5-LO-bzq_01v2 | LS | LOX5_HUMAN | † |
| LS4-09-5-LO-bzq_02v2 | LS | LOX5_HUMAN | † |
| LS4-09-5-LO-bzq_03v2 | LS | LOX5_HUMAN | † |
| LS4-09-5-LO-bzq-02-02-16-5LOX-benzoquinone_V2 | LS | LOX5_HUMAN | † |
| LS4-09-5-LO-zlt-20-01-16-zileuton | LS | LOX5_HUMAN | † |
| LS4-09-5-LO-zlt-20-01-16-zIleuton-derivatives | LS | LOX5_HUMAN | † |
| MR-2a3i-c0r-1.95-Z-1 | DS | MCR_HUMAN | (Praxmarer, 2011) |
| MR-2aa2-as4-1.95-Z-1 | DS | MCR_HUMAN | (Praxmarer, 2011) |
| MR-2aa2-as4-1.95-Z-2 | DS | MCR_HUMAN | (Praxmarer, 2011) |
| MR-2aa5-str-2.20-Z-1 | DS | MCR_HUMAN | (Praxmarer, 2011) |
| MR-2oax-snl-2.29-Z-1 | DS | MCR_HUMAN | (Praxmarer, 2011) |
| MR-Z001-1 | DS | MCR_HUMAN | (Praxmarer, 2011) |
| MR-Z001-2 | DS | MCR_HUMAN | (Praxmarer, 2011) |
| MR-Z002-1 | DS | MCR_HUMAN | (Praxmarer, 2011) |

| | | | |
|---|---|---|---|
| MR-Z002-2 | DS | MCR_HUMAN | (Praxmarer, 2011) |
| p38-MAPK-1a9u-sb2-800mod1_h | DS | MK14_HUMAN | (Humer, 2009) |
| p38-MAPK-1bl6-sb6-800mod1_h | DS | MK14_HUMAN | (Humer, 2009) |
| p38-MAPK-1bl7-sb4-800mod1_ha | DS | MK14_HUMAN | (Humer, 2009) |
| p38-MAPK-1bmk-sb5-800mod1_ha | DS | MK14_HUMAN | (Humer, 2009) |
| p38-MAPK-1ouk+shape | DS | MK14_HUMAN | (Humer, 2009) |
| p38-MAPK-1w82+shape | DS | MK14_HUMAN | (Humer, 2009) |
| p38-MAPK-1w83-1wbv+shape | DS | MK14_HUMAN | (Humer, 2009) |
| p38-MAPK-1wbvneu+shape | DS | MK14_HUMAN | (Humer, 2009) |
| p38-MAPK-2rg6-3cg2+shape | DS | MK14_HUMAN | (Humer, 2009) |
| p38-MAPK-3dt1-3ctq+shape | DS | MK14_HUMAN | (Humer, 2009) |
| NFkB-DNA-site-N003-1 | DS | NFKB1_HUMAN | * |
| LXR-1p8d | DS | NR1H2_HUMAN | (von Grafenstein et al., 2012) |
| LXR-1pq6 | DS | NR1H2_HUMAN | (von Grafenstein et al., 2012) |
| LXR-1pqc | DS | NR1H2_HUMAN | (von Grafenstein et al., 2012) |
| LXR-1uhl | DS | NR1H2_HUMAN | (von Grafenstein et al., 2012) |
| LXR-1upv | DS | NR1H2_HUMAN | (von Grafenstein et al., 2012) |
| LXR-1upw | DS | NR1H2_HUMAN | (von Grafenstein et al., 2012) |
| LXR-2acl | DS | NR1H2_HUMAN | (von Grafenstein et al., 2012) |
| LXR-3fal | DS | NR1H2_HUMAN | (von Grafenstein et al., 2012) |
| LXR-3fc6 | DS | NR1H2_HUMAN | (von Grafenstein et al., 2012) |
| FXR-1osh-fex-1.78-d-1 | DS | NR1H4_HUMAN | (Schuster et al., 2011b) |
| FXR-1osh-fex-1.78-d-1-s | DS | NR1H4_HUMAN | (Schuster et al., 2011b) |
| FXR-1osh-fex-1.78-d-2 | DS | NR1H4_HUMAN | (Schuster et al., 2011b) |
| FXR-1osh-fex-1.78-d-2-s | DS | NR1H4_HUMAN | (Schuster et al., 2011b) |
| FXR-3bej-muf-1.90-x-1 | DS | NR1H4_HUMAN | (Schuster et al., 2011b) |
| FXR-3bej-muf-1.90-x-1-s | DS | NR1H4_HUMAN | (Schuster et al., 2011b) |
| FXR-3bej-muf-1.90-x-2 | DS | NR1H4_HUMAN | (Schuster et al., 2011b) |
| FXR-3bej-muf-1.90-x-2-s | DS | NR1H4_HUMAN | (Schuster et al., 2011b) |
| FXR-3dct-o64-2.50-x-1 | DS | NR1H4_HUMAN | (Schuster et al., 2011b) |
| FXR-3dct-o64-2.50-x-1-s | DS | NR1H4_HUMAN | (Schuster et al., 2011b) |

| | | | |
|---|---|---|---|
| FXR-3dct-o64-2.50-x-2 | DS | NR1H4_HUMAN | (Schuster et al., 2011b) |
| FXR-3dct-o64-2.50-x-2-s | DS | NR1H4_HUMAN | (Schuster et al., 2011b) |
| FXR-3fli-33y-2.00-x-1 | DS | NR1H4_HUMAN | (Schuster et al., 2011b) |
| FXR-3fli-33y-2.00-x-1-s | DS | NR1H4_HUMAN | (Schuster et al., 2011b) |
| FXR-X008-1 | DS | NR1H4_HUMAN | (Schuster et al., 2011b) |
| cPLA2alpha-N002-2 | DS | PA2GA_HUMAN | (Noha et al., 2012) |
| COX-1-1cqe-flp-3.10-x-1 | DS | PGH1_HUMAN | (Temml et al., 2014) |
| COX-1-1pge-isf-3.50-x-2-s | DS | PGH1_HUMAN | (Temml et al., 2014) |
| COX-1-2ayl-flp-2.00-x-1 | DS | PGH1_HUMAN | (Temml et al., 2014) |
| COX-1-X017-1 | DS | PGH1_HUMAN | (Temml et al., 2014) |
| COX-1-1EQH4_LS3-01 | LS | PGH1_HUMAN | (Schuster et al., 2010) |
| COX-1-1PGE2_LS3-01 | LS | PGH1_HUMAN | (Schuster et al., 2010) |
| COX-1-1PGG2_LS3-01 | LS | PGH1_HUMAN | (Schuster et al., 2010) |
| COX-1-2AYL3_LS3-01 | LS | PGH1_HUMAN | (Schuster et al., 2010) |
| COX-1-2OYU2_LS3-01 | LS | PGH1_HUMAN | (Schuster et al., 2010) |
| COX-2-4cox-imn-2.90-x-2 | DS | PGH2_HUMAN | (Temml et al., 2014) |
| COX-2-6cox-s58-2.80-x-1-s | DS | PGH2_HUMAN | (Temml et al., 2014) |
| COX-2-3ln11_LS3-01 | LS | PGH2_HUMAN | (Schuster et al., 2010) |
| COX-2-3NTB1_LS3-01 | LS | PGH2_HUMAN | (Schuster et al., 2010) |
| COX-2-4COX2_LS3-01 | LS | PGH2_HUMAN | (Schuster et al., 2010) |
| COX-2-6COX3_LS3-01 | LS | PGH2_HUMAN | (Schuster et al., 2010) |
| PPARa-1i7g-az2-2.24-x-1 | DS | PPARA_HUMAN | (Markt, 2008) |
| PPARa-1i7g-az2-2.24-x-1-s | DS | PPARA_HUMAN | (Markt, 2008) |
| PPARa-1i7g-az2-2.24-x-2 | DS | PPARA_HUMAN | (Markt, 2008) |
| PPARa-1k7l-544-2.50-p-1 | DS | PPARA_HUMAN | (Markt, 2008) |
| PPARa-1k7l-544-2.50-p-1-s | DS | PPARA_HUMAN | (Markt, 2008) |
| PPARa-1k7l-544-2.50-p-2 | DS | PPARA_HUMAN | (Markt, 2008) |
| PPARa-1k7l-544-2.50-p-2-s | DS | PPARA_HUMAN | (Markt, 2008) |
| PPARa-1kkq-471-3.00-x-1 | DS | PPARA_HUMAN | (Markt, 2008) |
| PPARa-1kkq-471-3.00-x-1-s | DS | PPARA_HUMAN | (Markt, 2008) |
| PPARa-P007-1 | DS | PPARA_HUMAN | (Markt, 2008) |

| | | | |
|---|---|---|---|
| PPARd-1gwx-433-2.50-p-1 | DS | PPARD_HUMAN | (Markt, 2008) |
| PPARd-1gwx-433-2.50-p-1-s | DS | PPARD_HUMAN | (Markt, 2008) |
| PPARd-1y0s-331-2.65-p-1 | DS | PPARD_HUMAN | (Markt, 2008) |
| PPARd-1y0s-331-2.65-p-1-s | DS | PPARD_HUMAN | (Markt, 2008) |
| PPARd-1y0s-331-2.65-p-2-s | DS | PPARD_HUMAN | (Markt, 2008) |
| PPARd-1y0s-331-2.65-p-3 | DS | PPARD_HUMAN | (Markt, 2008) |
| PPARd-1y0s-331-2.65-p-3-s | DS | PPARD_HUMAN | (Markt, 2008) |
| PPARd-2awh-vca-2.00-p-1-s | DS | PPARD_HUMAN | (Markt, 2008) |
| PPARd-2awh-vca-2.00-p-2 | DS | PPARD_HUMAN | (Markt, 2008) |
| PPARd-2awh-vca-2.00-p-2-s | DS | PPARD_HUMAN | (Markt, 2008) |
| PPARd-2b50-vca-2.00-p-1-s | DS | PPARD_HUMAN | (Markt, 2008) |
| PPARd-2b50-vca-2.00-p-2 | DS | PPARD_HUMAN | (Markt, 2008) |
| PPARd-2b50-vca-2.00-p-2-s | DS | PPARD_HUMAN | (Markt, 2008) |
| PPARd-2baw-vca-2.30-p-1-s | DS | PPARD_HUMAN | (Markt, 2008) |
| PPARd-2j14-gni-2.80-p-1-s | DS | PPARD_HUMAN | (Markt, 2008) |
| PPARd-3gwx-epa-2.40-p-1-s | DS | PPARD_HUMAN | (Markt, 2008) |
| PPARd-3gwx-epa-2.40-p-2-s | DS | PPARD_HUMAN | (Markt, 2008) |
| PPARg-1fm6-brl-2.10-p-1 | DS | PPARG_HUMAN | (Markt, 2008) |
| PPARg-1fm6-brl-2.10-p-1-s | DS | PPARG_HUMAN | (Markt, 2008) |
| PPARg-1fm9-570-2.10-x-1 | DS | PPARG_HUMAN | (Markt, 2008) |
| PPARg-1fm9-570-2.10-x-1-s | DS | PPARG_HUMAN | (Markt, 2008) |
| PPARg-1fm9-570-2.10-x-2 | DS | PPARG_HUMAN | (Markt, 2008) |
| PPARg-1fm9-570-2.10-x-2-s | DS | PPARG_HUMAN | (Markt, 2008) |
| PPARg-1i7i-az2-2.35-x-1 | DS | PPARG_HUMAN | (Markt, 2008) |
| PPARg-1i7i-az2-2.35-x-1-s | DS | PPARG_HUMAN | (Markt, 2008) |
| PPARg-1k74-544-2.30-p-1-s | DS | PPARG_HUMAN | (Markt, 2008) |
| PPARg-1knu-ypa-2.50-p-1-s | DS | PPARG_HUMAN | (Markt, 2008) |
| PPARg-1knu-ypa-2.50-p-2-s | DS | PPARG_HUMAN | (Markt, 2008) |
| PPARg-1nyx-drf-2.65-p-1 | DS | PPARG_HUMAN | (Markt, 2008) |
| PPARg-1nyx-drf-2.65-p-1-s | DS | PPARG_HUMAN | (Markt, 2008) |
| PPARg-1nyx-drf-2.65-p-2-s | DS | PPARG_HUMAN | (Markt, 2008) |

| | | | |
|---|---|---|---|
| PPARg-1wm0-plb-2.90-p-1-s | DS | PPARG_HUMAN | (Markt, 2008) |
| PPARg-1zeo-c01-2.50-p-1 | DS | PPARG_HUMAN | (Markt, 2008) |
| PPARg-1zeo-c01-2.50-p-1-s | DS | PPARG_HUMAN | (Markt, 2008) |
| PPARg-1zgy-brl-1.80-p-1-s | DS | PPARG_HUMAN | (Markt, 2008) |
| PPARg-2ath-3ea-2.28-p-1-s | DS | PPARG_HUMAN | (Markt, 2008) |
| PPARg-2f4b-eha-2.07-p-1-s | DS | PPARG_HUMAN | (Markt, 2008) |
| PPARg-2fvj-ro0-1.99-p-1-s | DS | PPARG_HUMAN | (Markt, 2008) |
| PPARg-2g0g-sp0-2.54-p-1 | DS | PPARG_HUMAN | (Markt, 2008) |
| PPARg-2g0g-sp0-2.54-p-1-s | DS | PPARG_HUMAN | (Markt, 2008) |
| PPARg-2g0h-sp3-2.30-p-1-s | DS | PPARG_HUMAN | (Markt, 2008) |
| PPARg-2gtk-208-2.10-p-1-s | DS | PPARG_HUMAN | (Markt, 2008) |
| PPARg-2hfp-nsi-2.00-p-1-s | DS | PPARG_HUMAN | (Markt, 2008) |
| PPARg-2prg-brl-2.30-p-1 | DS | PPARG_HUMAN | (Markt, 2008) |
| PPARg-2prg-brl-2.30-p-1-s | DS | PPARG_HUMAN | (Markt, 2008) |
| PPARg-2q59-240-2.20-p-1-s | DS | PPARG_HUMAN | (Markt, 2008) |
| PPARg-4prg-072-2.90-p-1-s | DS | PPARG_HUMAN | (Markt, 2008) |
| PPARg-P002-1 | DS | PPARG_HUMAN | (Markt, 2008) |
| mPGES-1-Hypo_62_01-non-acidic | DS | PTGES_HUMAN | (Noha et al., 2015) |
| mPGES-1-X019-1 | DS | PTGES_HUMAN | (Noha et al., 2015) |
| mPGES-1-X019-2 | DS | PTGES_HUMAN | (Noha et al., 2015) |
| mPGES-1-model-LS3-01 | LS | PTGES_HUMAN | (Noha et al., 2015) |
| PTP1b-1bzc-tpi-2.35-d-2 | DS | PTN1_HUMAN | (Herdlinger, 2016) |
| PTP1b-1bzh-flt-2.10-d-1 | DS | PTN1_HUMAN | (Herdlinger, 2016) |
| PTP1b-1bzj-pic-2.25-d-1 | DS | PTN1_HUMAN | (Herdlinger, 2016) |
| PTP1b-1c83-oai-1.80-d-1-s | DS | PTN1_HUMAN | (Herdlinger, 2016) |
| PTP1b-1c83-oai-1.80-d-2 | DS | PTN1_HUMAN | (Herdlinger, 2016) |
| PTP1b-1c83-oai-1.80-d-2-s | DS | PTN1_HUMAN | (Herdlinger, 2016) |
| PTP1b-1c84-761-2.35-d-1 | DS | PTN1_HUMAN | (Herdlinger, 2016) |
| PTP1b-1c84-761-2.35-d-1-s | DS | PTN1_HUMAN | (Herdlinger, 2016) |
| PTP1b-1c84-761-2.35-d-2 | DS | PTN1_HUMAN | (Herdlinger, 2016) |
| PTP1b-1c84-761-2.35-d-2-s | DS | PTN1_HUMAN | (Herdlinger, 2016) |

| | | | |
|---|---|---|---|
| PTP1b-1c84-761-2.35-d-3 | DS | PTN1_HUMAN | (Herdlinger, 2016) |
| PTP1b-1c84-761-2.35-d-3-s | DS | PTN1_HUMAN | (Herdlinger, 2016) |
| PTP1b-1c85-oba-2.72-d-1 | DS | PTN1_HUMAN | (Herdlinger, 2016) |
| PTP1b-1c85-oba-2.72-d-1-s | DS | PTN1_HUMAN | (Herdlinger, 2016) |
| PTP1b-1c86-opa-2.30-d-1 | DS | PTN1_HUMAN | (Herdlinger, 2016) |
| PTP1b-1c86-opa-2.30-d-1-s | DS | PTN1_HUMAN | (Herdlinger, 2016) |
| PTP1b-1c87-opa-2.10-d-1 | DS | PTN1_HUMAN | (Herdlinger, 2016) |
| PTP1b-1c87-opa-2.10-d-1-s | DS | PTN1_HUMAN | (Herdlinger, 2016) |
| PTP1b-1c88-ota-1.80-d-1 | DS | PTN1_HUMAN | (Herdlinger, 2016) |
| PTP1b-1ecv-878-1.95-d-1 | DS | PTN1_HUMAN | (Herdlinger, 2016) |
| PTP1b-1ecv-878-1.95-d-1-s | DS | PTN1_HUMAN | (Herdlinger, 2016) |
| PTP1b-1ecv-878-1.95-d-2 | DS | PTN1_HUMAN | (Herdlinger, 2016) |
| PTP1b-1ecv-878-1.95-d-2-s | DS | PTN1_HUMAN | (Herdlinger, 2016) |
| PTP1b-1ecv-878-1.95-d-3 | DS | PTN1_HUMAN | (Herdlinger, 2016) |
| PTP1b-1ecv-878-1.95-d-3-s | DS | PTN1_HUMAN | (Herdlinger, 2016) |
| PTP1b-1g7g-inx-2.20-d-1 | DS | PTN1_HUMAN | (Herdlinger, 2016) |
| PTP1b-1gfy-col-2.13-d-1 | DS | PTN1_HUMAN | (Herdlinger, 2016) |
| PTP1b-1gfy-col-2.13-d-1-s | DS | PTN1_HUMAN | (Herdlinger, 2016) |
| PTP1b-1jf7-tbh-2.20-d-1 | DS | PTN1_HUMAN | (Herdlinger, 2016) |
| PTP1b-1kak-fnp-2.50-d-1 | DS | PTN1_HUMAN | (Herdlinger, 2016) |
| PTP1b-1kak-fnp-2.50-d-2 | DS | PTN1_HUMAN | (Herdlinger, 2016) |
| PTP1b-1l8g-dbd-2.50-d-1 | DS | PTN1_HUMAN | (Herdlinger, 2016) |
| PTP1b-1nl9-989-2.40-d-1 | DS | PTN1_HUMAN | (Herdlinger, 2016) |
| PTP1b-1nl9-989-2.40-d-1-s | DS | PTN1_HUMAN | (Herdlinger, 2016) |
| PTP1b-1nny-515-2.40-d-1 | DS | PTN1_HUMAN | (Herdlinger, 2016) |
| PTP1b-1no6-794-2.40-d-1 | DS | PTN1_HUMAN | (Herdlinger, 2016) |
| PTP1b-1no6-794-2.40-d-1-s | DS | PTN1_HUMAN | (Herdlinger, 2016) |
| PTP1b-1nwl-964-2.40-d-1 | DS | PTN1_HUMAN | (Herdlinger, 2016) |
| PTP1b-1nwl-964-2.40-d-1-s | DS | PTN1_HUMAN | (Herdlinger, 2016) |
| PTP1b-1nz7-901-2.40-d-1 | DS | PTN1_HUMAN | (Herdlinger, 2016) |
| PTP1b-1ony-588-2.15-d-1 | DS | PTN1_HUMAN | (Herdlinger, 2016) |

| | | | |
|---|---|---|---|
| PTP1b-1ony-588-2.15-d-1-s | DS | PTN1_HUMAN | (Herdlinger, 2016) |
| PTP1b-1onz-968-2.40-d-1 | DS | PTN1_HUMAN | (Herdlinger, 2016) |
| PTP1b-1onz-968-2.40-d-1-s | DS | PTN1_HUMAN | (Herdlinger, 2016) |
| PTP1b-1ph0-418-2.20-d-1 | DS | PTN1_HUMAN | (Herdlinger, 2016) |
| PTP1b-1ph0-418-2.20-d-2 | DS | PTN1_HUMAN | (Herdlinger, 2016) |
| PTP1b-1ph0-418-2.20-d-2-s | DS | PTN1_HUMAN | (Herdlinger, 2016) |
| PTP1b-1pxh-sna-2.15-d-1 | DS | PTN1_HUMAN | (Herdlinger, 2016) |
| PTP1b-1pyn-941-2.20-d-1 | DS | PTN1_HUMAN | (Herdlinger, 2016) |
| PTP1b-1q1m-234-2.60-d-1 | DS | PTN1_HUMAN | (Herdlinger, 2016) |
| PTP1b-1q6j-335-2.20-d-1 | DS | PTN1_HUMAN | (Herdlinger, 2016) |
| PTP1b-1q6j-335-2.20-d-1-s | DS | PTN1_HUMAN | (Herdlinger, 2016) |
| PTP1b-1q6m-p27-2.20-d-1 | DS | PTN1_HUMAN | (Herdlinger, 2016) |
| PTP1b-1q6m-p27-2.20-d-1-s | DS | PTN1_HUMAN | (Herdlinger, 2016) |
| PTP1b-1q6n-p90-2.10-d-1 | DS | PTN1_HUMAN | (Herdlinger, 2016) |
| PTP1b-1q6n-p90-2.10-d-1-s | DS | PTN1_HUMAN | (Herdlinger, 2016) |
| PTP1b-1q6p-213-2.30-d-1 | DS | PTN1_HUMAN | (Herdlinger, 2016) |
| PTP1b-1q6s-214-2.20-d-1 | DS | PTN1_HUMAN | (Herdlinger, 2016) |
| PTP1b-1q6s-214-2.20-d-1-s | DS | PTN1_HUMAN | (Herdlinger, 2016) |
| PTP1b-1q6t-600-2.30-d-1 | DS | PTN1_HUMAN | (Herdlinger, 2016) |
| PTP1b-1qxk-429-2.30-d-1 | DS | PTN1_HUMAN | (Herdlinger, 2016) |
| PTP1b-2bgd-t1d-2.40-d-1 | DS | PTN1_HUMAN | (Herdlinger, 2016) |
| PTP1b-2cne-dfj-1.80-d-1 | DS | PTN1_HUMAN | (Herdlinger, 2016) |
| PTP1b-2cne-dfj-1.80-d-1-s | DS | PTN1_HUMAN | (Herdlinger, 2016) |
| PTP1b-2cne-dfj-1.80-d-2 | DS | PTN1_HUMAN | (Herdlinger, 2016) |
| PTP1b-2cne-dfj-1.80-d-2-s | DS | PTN1_HUMAN | (Herdlinger, 2016) |
| PTP1b-2cnf-f32-2.20-d-1 | DS | PTN1_HUMAN | (Herdlinger, 2016) |
| PTP1b-2cnf-f32-2.20-d-1-s | DS | PTN1_HUMAN | (Herdlinger, 2016) |
| PTP1b-2cnf-f32-2.20-d-2 | DS | PTN1_HUMAN | (Herdlinger, 2016) |
| PTP1b-2cnf-f32-2.20-d-2-s | DS | PTN1_HUMAN | (Herdlinger, 2016) |
| PTP1b-2cng-ize-1.90-d-1 | DS | PTN1_HUMAN | (Herdlinger, 2016) |
| PTP1b-2cng-ize-1.90-d-1-s | DS | PTN1_HUMAN | (Herdlinger, 2016) |

| | | | |
|---|---|---|---|
| PTP1b-2cnh-izb-1.80-d-1 | DS | PTN1_HUMAN | (Herdlinger, 2016) |
| PTP1b-2cnh-izb-1.80-d-1-s | DS | PTN1_HUMAN | (Herdlinger, 2016) |
| PTP1b-2cni-izf-2.00-d-1 | DS | PTN1_HUMAN | (Herdlinger, 2016) |
| PTP1b-2cni-izf-2.00-d-1-s | DS | PTN1_HUMAN | (Herdlinger, 2016) |
| PTP2b-1g7f-inz-1.80-d-1 | DS | PTN1_HUMAN | (Herdlinger, 2016) |
| LS3-03b-1C85-1-model | LS | PTN1_HUMAN | (Herdlinger, 2016) |
| LS3-03b-1C85-2-model | LS | PTN1_HUMAN | (Herdlinger, 2016) |
| LS3-03b-1PH0-model | LS | PTN1_HUMAN | (Herdlinger, 2016) |
| LS3-03b-1PYN-model | LS | PTN1_HUMAN | (Herdlinger, 2016) |
| LS3-03b-1Q6S-model | LS | PTN1_HUMAN | (Herdlinger, 2016) |
| LS3-03b-1T49-model | LS | PTN1_HUMAN | (Herdlinger, 2016) |
| LS3-03b-2CM7-model | LS | PTN1_HUMAN | (Herdlinger, 2016) |
| LS3-03b-2CNG-model | LS | PTN1_HUMAN | (Herdlinger, 2016) |
| LS3-03b-2F71-model | LS | PTN1_HUMAN | (Herdlinger, 2016) |
| LS3-03b-2QBS-model | LS | PTN1_HUMAN | (Herdlinger, 2016) |
| IKK2-3rzf-xnm-4.00-n-1 | DS | Q6INT1_XENLA | (Noha et al., 2011) |
| IKK2-N001-6 | DS | Q6INT1_XENLA | (Noha et al., 2011) |
| 5a-red-R003-2 | DS | S5A2_HUMAN | (Rhöse, 2010) |
| 5a-red-R003-4 | DS | S5A2_HUMAN | (Rhöse, 2010) |
| STS-LS3.12-model10-mostafa-2012a-21-maltais-2009a-55-XVols-binding-site | LS | STS_HUMAN | (Grienke et al., 2020) |
| STS-LS3.12-model11-lehr-2005a-4f-nussbaumer-2003b-6g | LS | STS_HUMAN | (Grienke et al., 2020) |
| STS-LS3.12-model12-mostafa-2012a-21-maltais-2009a-55-XVols-binding-site-HBD | LS | STS_HUMAN | (Grienke et al., 2020) |

* not published. † in preparation.

### S-2 PIPELINE PILOT PROTOCOL:

ChemDraw's native file format '.cdx' assignes stereochemistry, which can be preserved when converted using this PipelinePilot protocol called 'chemdraw_to_sd_smiles_fabian.xml'. The removal of duplicate entries is necessary, since various names of the compounds can be found in literature, and consequently in the hand-drawn library. The protocol is further provided on GitHub (https://github.com/fmayr/DHC_TargetPrediction).

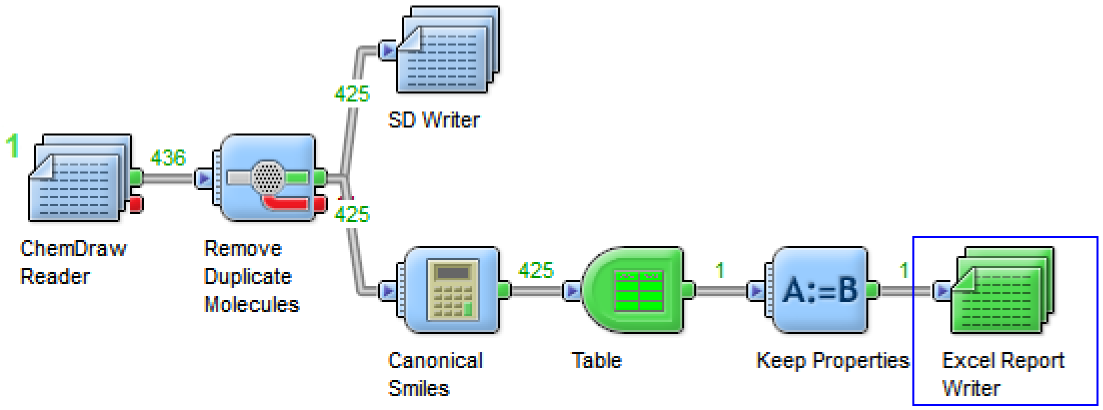


Figure S 2. Pipeline Pilot protocol 'chemdraw_to_sd_smiles_fabian.xml' used in this study.

### S-3 FILE SCHEME:

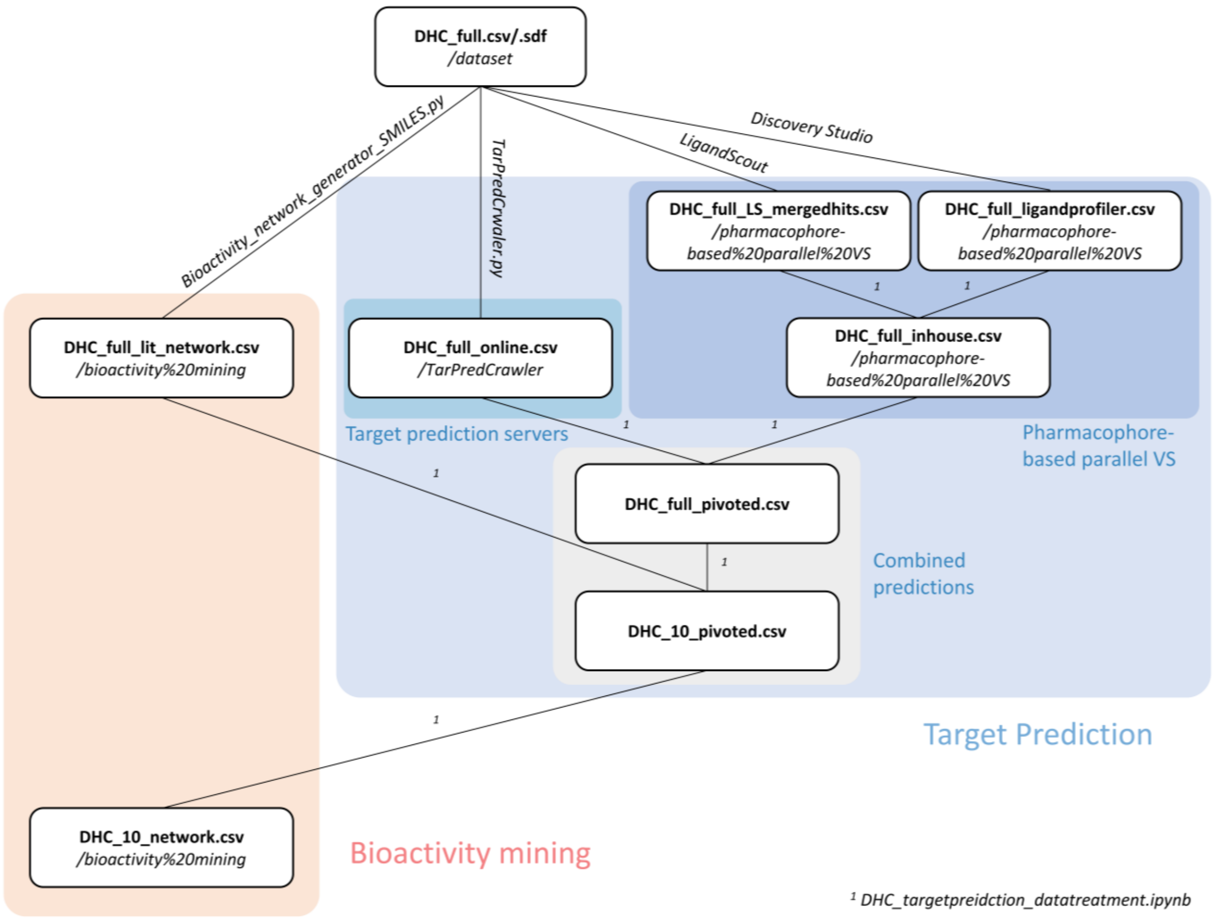


Figure S 3. File scheme describing the course of this study. All files are available for download on GitHub (https://github.com/fmayr/DHC_TargetPrediction), relative paths are written in italics.

### S-4 LIGAND PROFILER PARAMETER SETTINGS

Table S 4. Parameter settings used for the Ligand Profiler protocol used in parallel VS with Discovery Studio.

| **Protocol Settings** | **Protocol.pr_xml** | |
|---|---|---|
| | **Input Ligands** | DHC_full.sdf |
| | **Input LigandProfilerDB Pharmacophores** | |
| | **Input PharmaDB Pharmacophores** | |
| **Model Selection** | Most Selective | |
| | **Conformation Generation** | BEST |
| **Maximum Conformations** | 255 | |
| **Discard Existing Conformations** | WAHR | |
| **Energy Threshold** | 20 | |
| **Ring Fragments File** | | |
| **Save Conformations** | FALSCH | |
| | **Advanced** | |
| **Input Type** | Ligands | |
| **Input Database** | Sample | |
| **Input Database Limit Hits** | First N | |
| **Input Database Maximum** | 300 | |
| **Input Database Hitlist** | | |
| **Fitting Method** | Rigid | |
| **Maximum Omitted Features** | 0 | |
| **Minimum Interfeature Distance** | 0.00001 | |
| **Scale Fit Values** | WAHR | |
| **Prune Empty Fits** | WAHR | |
| **Prune Missed Molecules** | WAHR | |
| **Keep Input Conformations** | FALSCH | |
| **Save Aligned Ligands** | FALSCH | |
| **Activity Property** | | |
| **Catalyst Parameter File** | | |
| | **Parallel Processing** | FALSCH |

### S-5 QUALITY CONTROL OF 1 – 10:

#### Phloretin (1) CAS: 60-82-2

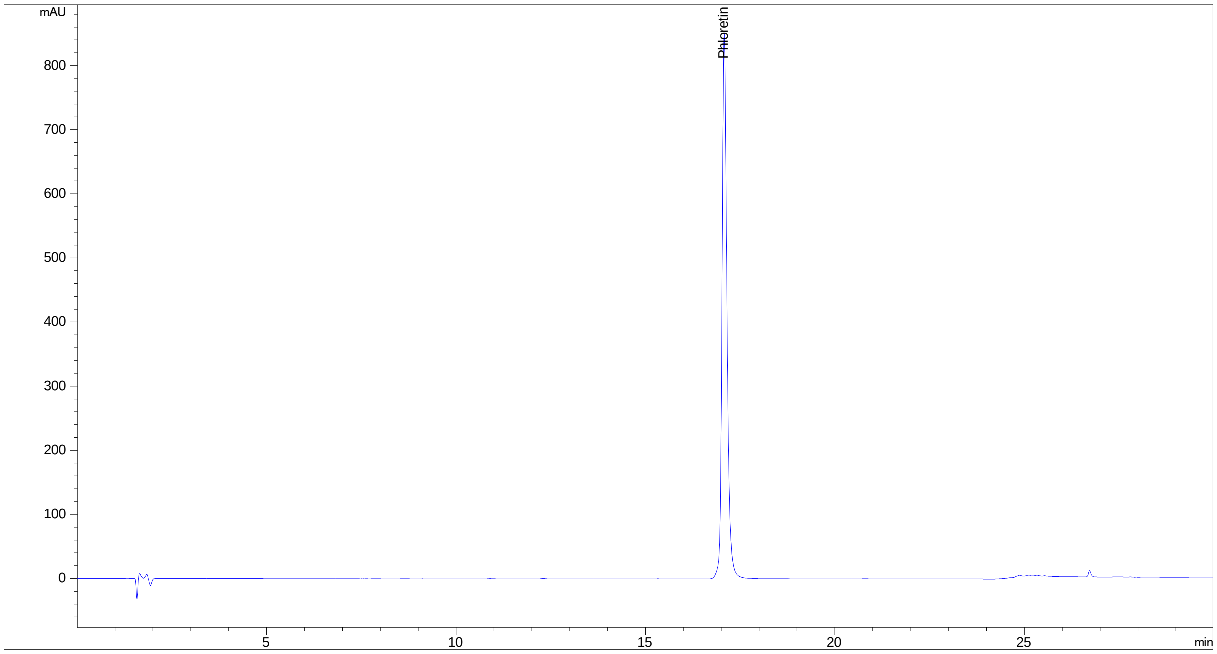


#### 3-OH-phloretin (2) CAS: 57765-66-9

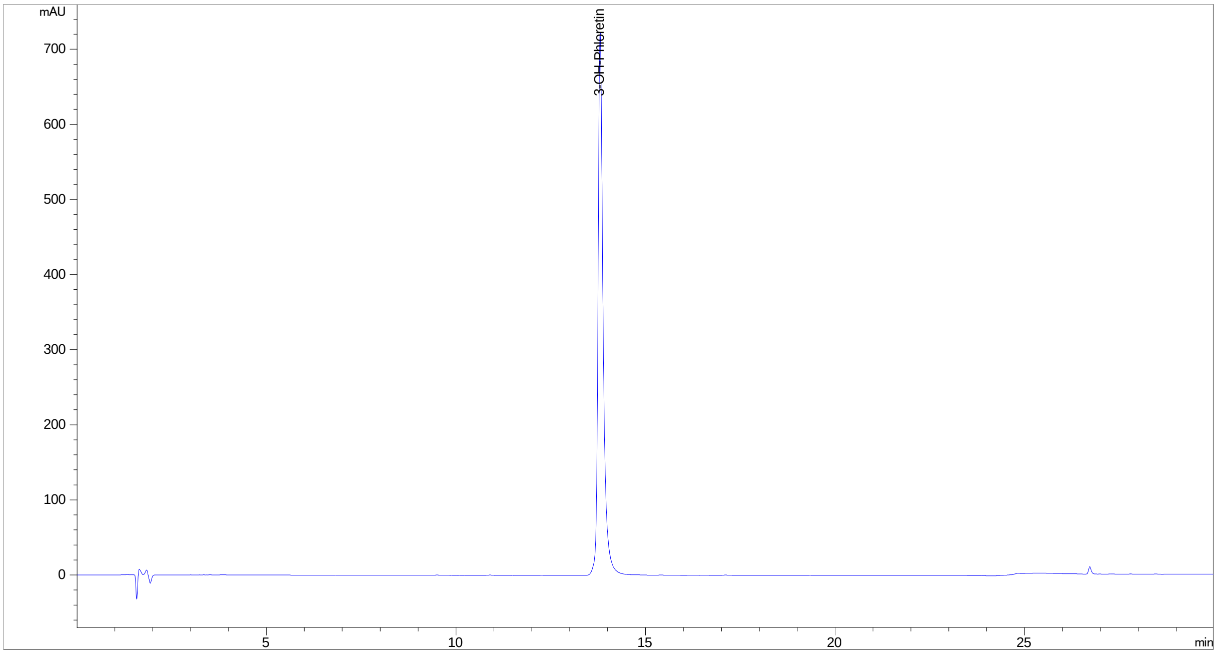

#### 2',6'-dihydroxy-4'-methoxy DHC (3) CAS: 35241-55-5

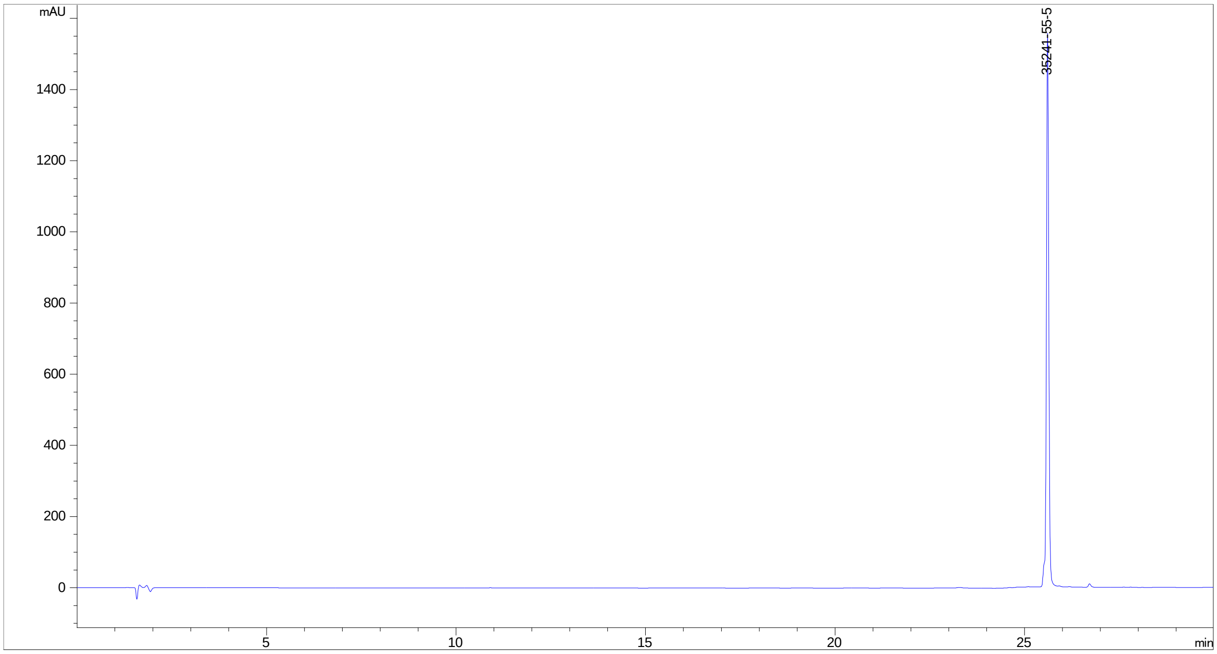


#### Asebogenin (4) CAS: 35241-54-4

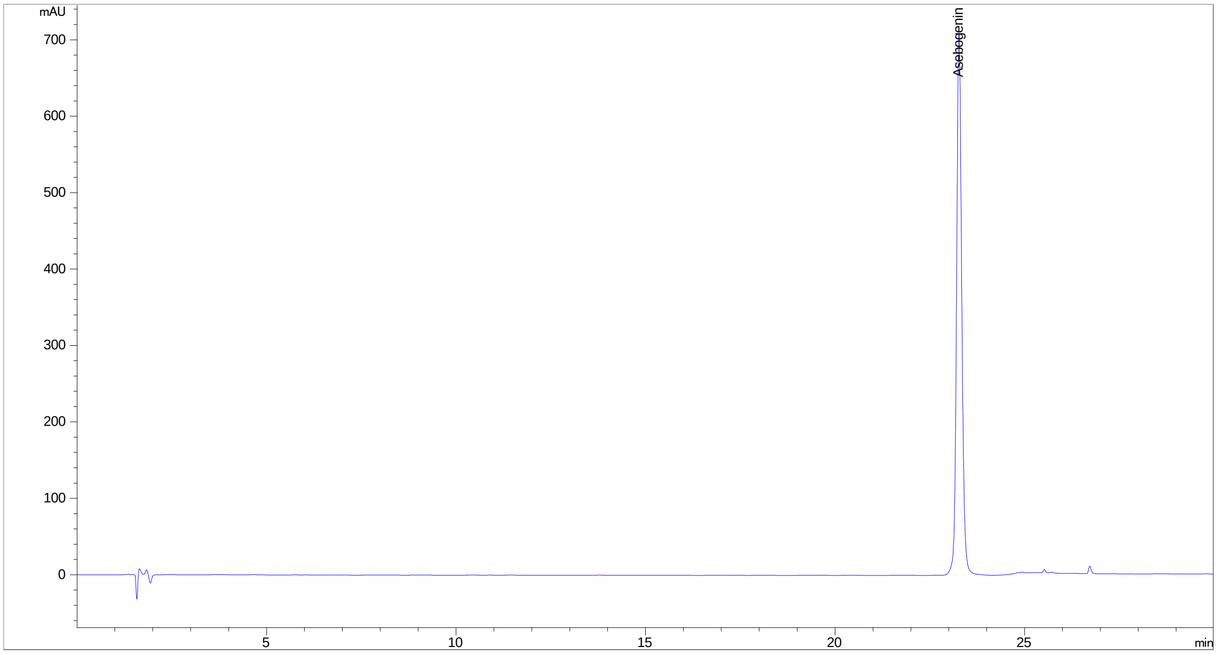

#### Calomelanen (5) CAS: 520-42-3

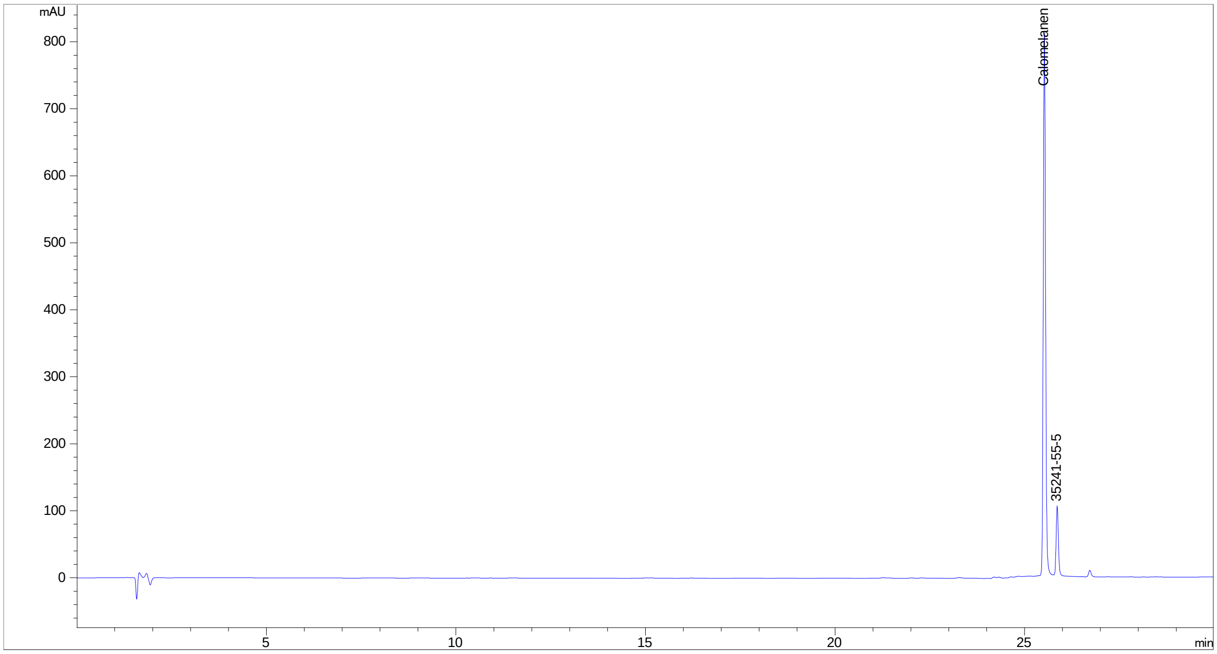


#### Sieboldin (6) CAS: 18777-73-6

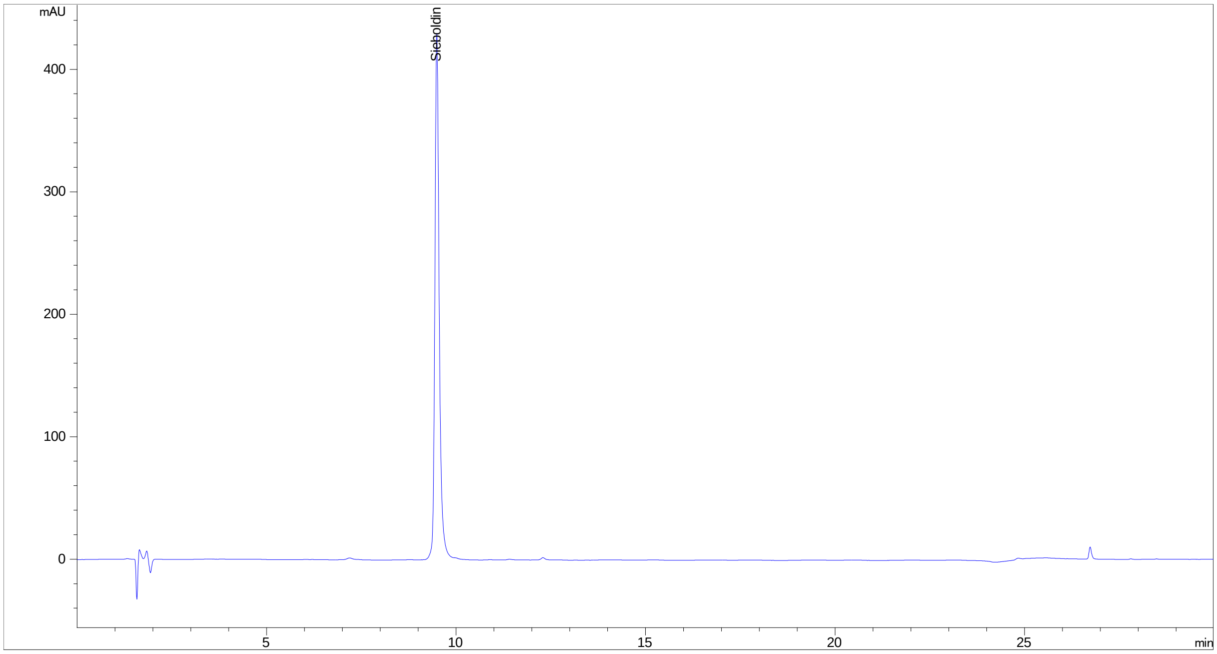

#### Phloridzin (7) CAS: 60-81-1

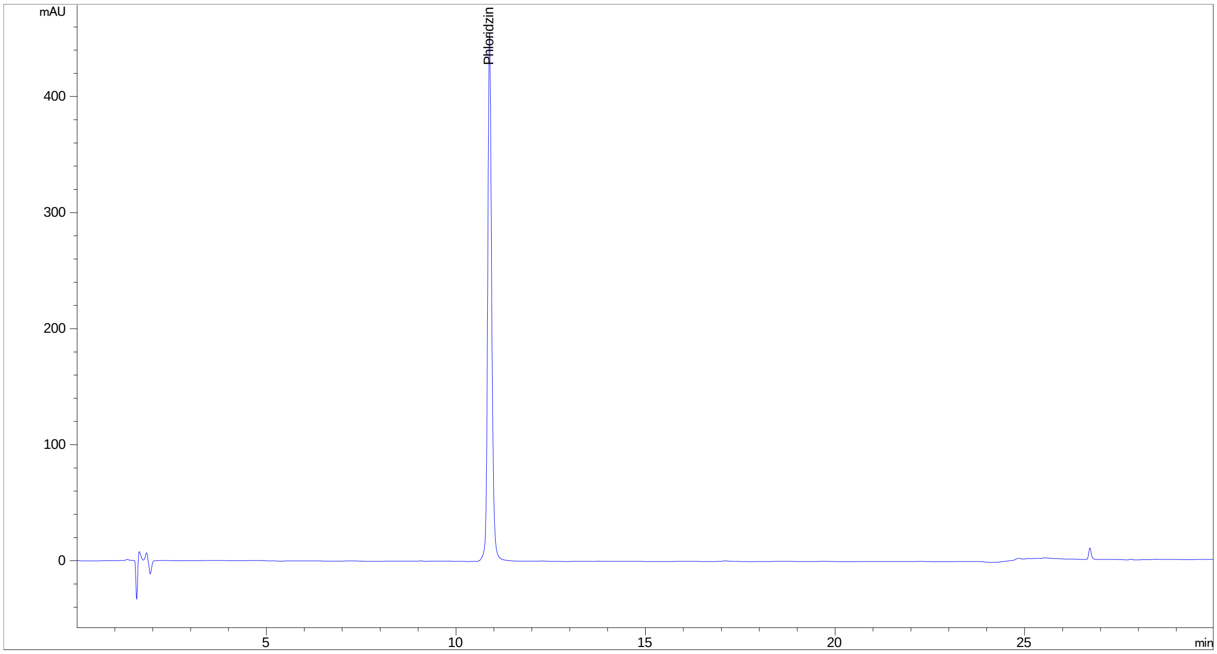


#### Trilobatin (8) CAS: 4192-90-9

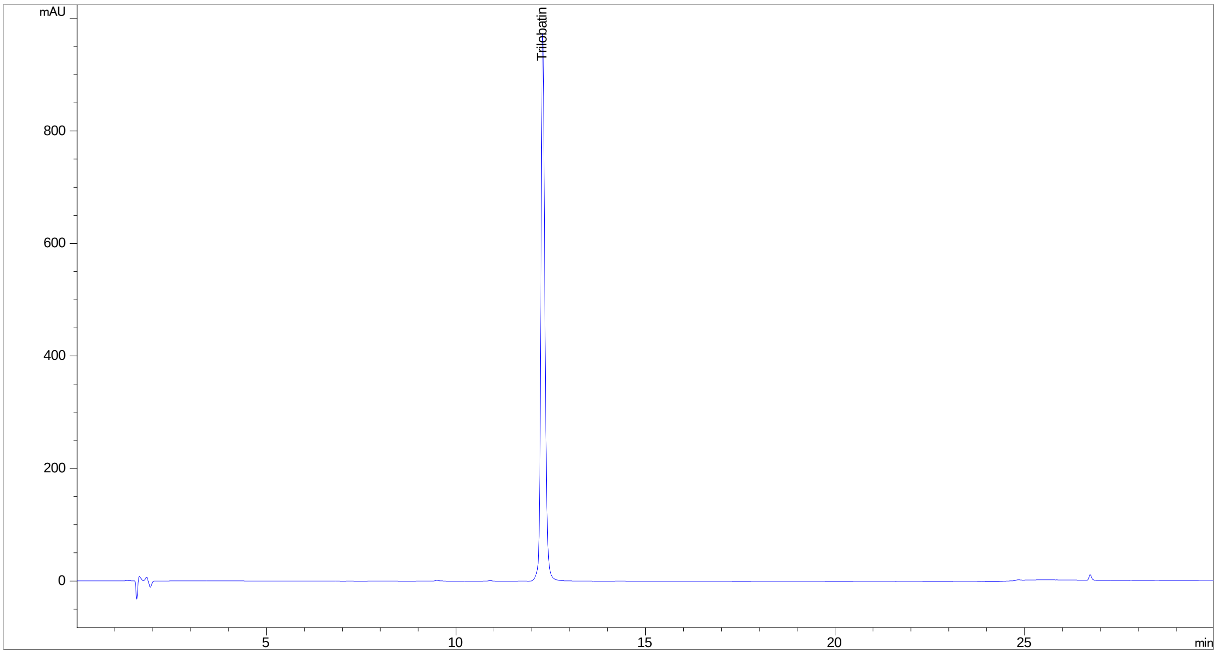

**Phloretin-2'-xyloglucoside (9) CAS: 145758-09-4**

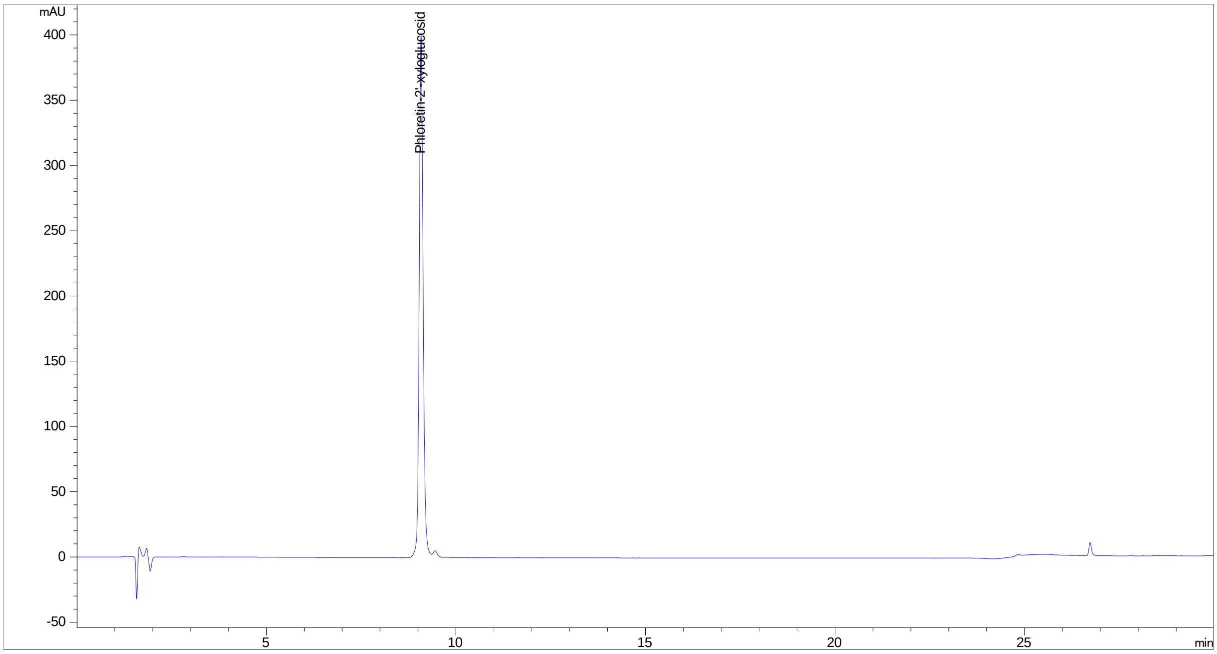


**Neohesperidin DHC (10) CAS: 20702-77-6**

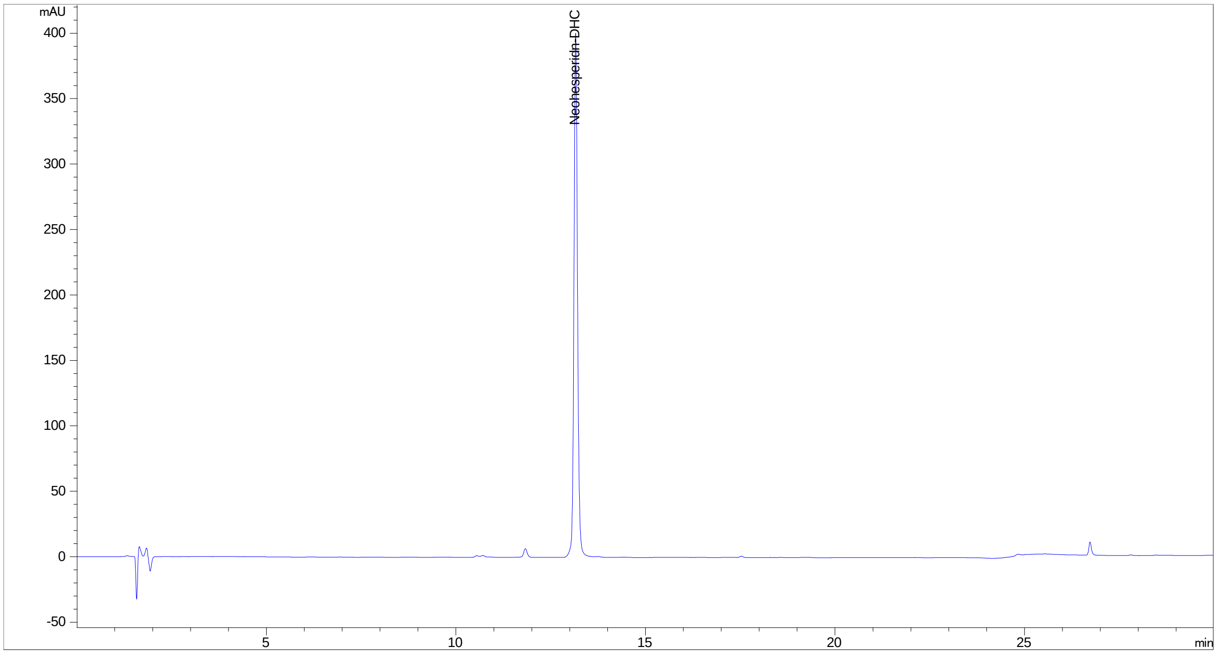

### S-6 POSITIVE CONTROLS:

| Assay | Name | CAS | Structure | Reference |
|---|---|---|---|---|
| **3β HSD1** | Trilostane | 13647-35-3 | 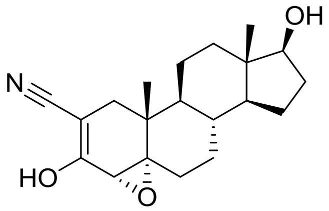 | (Samandari et al., 2007; Udhane et al., 2017) |
| **11β HSD1** **11β HSD2** | 18β-Glycyrrhetinic acid | 471-53-4 | 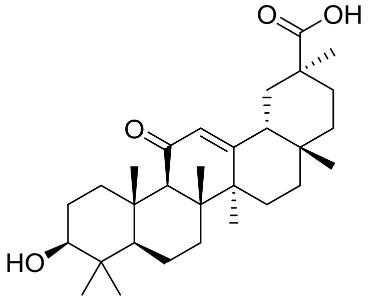 | (D. V. Kratschmar et al., 2011b) |
| **17β HSD2** **17β HSD4** | compound 19 | 1340482-23-6 | 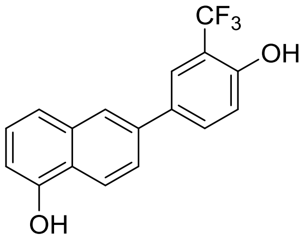 | (Wetzel et al., 2011) |
| **17β HSD3** | compound 24 | 873206-61-2 | 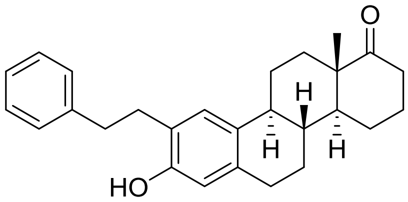 | (Möller et al., 2009) |
| **5-LO** | Zileuton | 111406-87-2 | 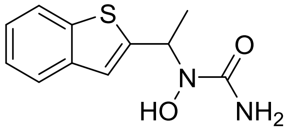 | (Koeberle et al., 2008) |
| **aromatase** | anastrozole | 120511-73-1 | 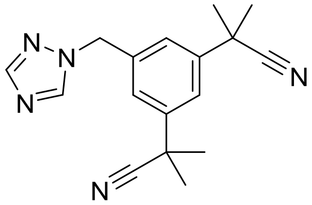 | (Pandey et al., 2007) |

| | | | | |
|---|---|---|---|---|
| **AKR1C3** | compound 2-9 | 745028-76-6 | 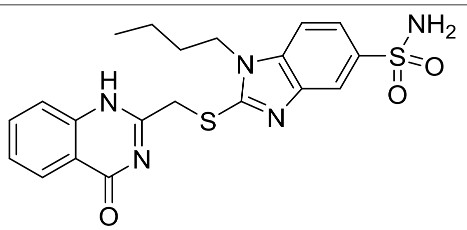 | (Schuster et al., 2011a) |
| **COX-1** | Indomethacin | 53-86-1 | 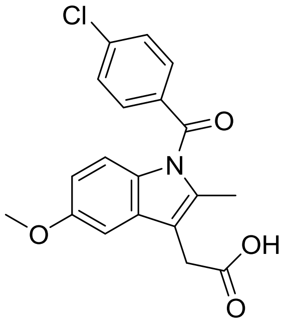 | (Schaible et al., 2014) |
| **CYP17A1** | Abiraterone | 154229-19-3 | 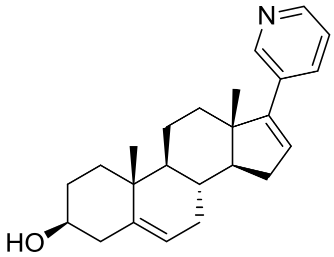 | (Samandari et al., 2007; Udhane et al., 2017) |
| **sEH** | AUDA | 479413-70-2 | 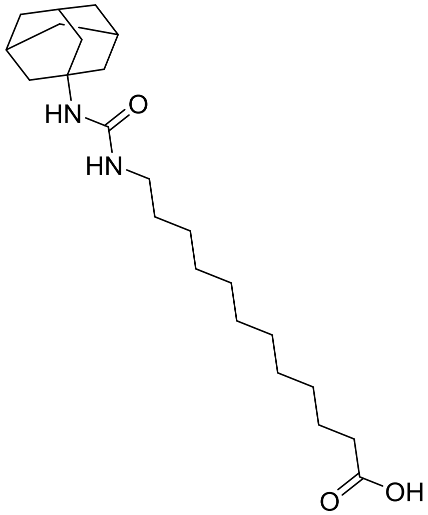 | (Waltenberger et al., 2016) |

### S-7 REFERENCES:


Akram, M., Patt, M., Kaserer, T., Temml, V., Waratchareeyakul, W., Kratschmar, D. V., . . . Schuster, D. (2019). Identification of the fungicide epoxiconazole by virtual screening and biological assessment as inhibitor of human 11β-hydroxylase and aldosterone synthase. *J.*

*Steroid Biochem. Mol. Biol., 192*, 105358. doi: https://doi.org/10.1016/j.jsbmb.2019.04.007

Brunner, F. (2011). *A Pharmacophore model collection for estrogen receptor ligands using ligandscout.* (Mag. pharm.), Leopold-Franzens Universität Innsbruck, Innsbruck. Retrieved from https://permalink.obvsg.at/AC08718721

Fiala, J. (2017). *Pharmacophore modeling and virtual screening for 17β -hydroxysteroid dehydrogenase 7 inhibitors.* (Mag. pharm.), Leopold-Franzens Universität Innsbruck, Innsbruck. Retrieved from https://permalink.obvsg.at/AC13701540

Grienke, U., Kaserer, T., Kirchweger, B., Lambrinidis, G., Kandel, R. T., Foster, P. A., . . . Rollinger, J. M. (2020). Steroid sulfatase inhibiting lanostane triterpenes – Structure activity relationship and in silico insights. *Bioorg. Chem., 95*, 103495. doi: https://doi.org/10.1016/j.bioorg.2019.103495

Herdlinger, S. (2016). *Computer-assisted structure-activity studies on protein tyrosine phosphatase 1B (PTP1B)-inhibitors.* (Mag. pharm.), Leopold-Franzens Universität Innsbruck, Innsbruck. Retrieved from https://permalink.obvsg.at/AC13481365

Hochleitner, J., Akram, M., Ueberall, M., Davis, R. A., Waltenberger, B., Stuppner, H., . . . Schuster, D. (2017). A combinatorial approach for the discovery of cytochrome P450 2D6 inhibitors from nature. *Sci. Rep., 7*(1), 8071. doi: 10.1038/s41598-017-08404-0

Humer, K. (2009). *3D - pharmacophore modeling for kinases involved in inflammation.* (Mag. pharm.), Leopold-Franzens Universität Innsbruck, Innsbruck. Retrieved from https://bibsearch.uibk.ac.at/AC07878262

Koeberle, A., Siemoneit, U., Bühring, U., Northoff, H., Laufer, S., Albrecht, W., & Werz, O. (2008). Licofelone suppresses prostaglandin $E_2$ formation by interference with the inducible microsomal prostaglandin $E_2$ synthase-1. *J. Pharmacol. Exp. Ther., 326*(3), 975. doi: 10.1124/jpet.108.139444

Kratschmar, D. V., Vuorinen, A., Da Cunha, T., Wolber, G., Classen-Houben, D., Doblhoff, O., . . . Odermatt, A. (2011a). Characterization of activity and binding mode of glycyrrhetinic acid derivatives inhibiting 11β-hydroxysteroid dehydrogenase type 2. *The Journal of Steroid Biochemistry and Molecular Biology, 125*(1), 129-142. doi: https://doi.org/10.1016/j.jsbmb.2010.12.019

Kratschmar, D. V., Vuorinen, A., Da Cunha, T., Wolber, G., Classen-Houben, D., Doblhoff, O., . . . Odermatt, A. (2011b). Characterization of activity and binding mode of glycyrrhetinic acid derivatives inhibiting 11β-hydroxysteroid dehydrogenase type 2. *J. Steroid Biochem. Mol. Biol.*, *125*(1), 129-142. doi: https://doi.org/10.1016/j.jsbmb.2010.12.019

Kurzemann, F. (2016). *Pharmacophore modeling and virtual screening for 3β -hydroxysteroid-dehydrogenase inhibitors.* (Mag. pharm.), Innsbruck, Leopold-Franzens Universität Innsbruck. Retrieved from https://permalink.obvsg.at/AC13434797

Linder, T. (2010). *Pharmacophore modeling and virtual screening for glucocorticoid receptor ligands: unraveling anti-inflammatory or endocrine disrupting effects of chemicals.* (Mag. pharm.), Leopold-Franzens Universität Innsbruck, Innsbruck. Retrieved from https://permalink.obvsg.at/AC08358721

Markt, P. (2008). *3D Virtual high-throughput screening : discovery of novel lead structures and validation of the parallel screening approach with focus on targets involved in the metabolic syndrome . Kapitel 3 auch erschienen in: Journal of Computer-aided Molecular Design, 2008, online, doi: 10.1007/s10822-007-9163-6, als Artikel sobald wie möglich / Kapitel 4 auch erschienen in: Journal of Computer-aided Molecular Design, 2007, Band 10-11, Seiten 575-590.* (Dr. rer. nat.), Leopold-Franzens Universität Innsbruck, Innsbruck.

Möller, G., Deluca, D., Gege, C., Rosinus, A., Kowalik, D., Peters, O., . . . Hillisch, A. (2009). Structure-based design, synthesis and in vitro characterization of potent 17β-hydroxysteroid dehydrogenase type 1 inhibitors based on 2-substitutions of estrone and D-homo-estrone. *Bioorg. Med. Chem. Lett.*, *19*(23), 6740-6744. doi: https://doi.org/10.1016/j.bmcl.2009.09.113

Noha, S. M. (2013). *Application of 3D virtual screening techniques for the identification of novel anti-inflammatory plant constituents.* (PhD), Leopold-Franzens Universität Innsbruck, Innsbruck. Retrieved from https://permalink.obvsg.at/AC10774943

Noha, S. M., Atanasov, A. G., Schuster, D., Markt, P., Fakhrudin, N., Heiss, E. H., . . . Wolber, G. (2011). Discovery of a novel IKK-β inhibitor by ligand-based virtual screening techniques. *Bioorg. Med. Chem. Lett.*, *21*(1), 577-583. doi: https://doi.org/10.1016/j.bmcl.2010.10.051

Noha, S. M., Fischer, K., Koeberle, A., Garscha, U., Werz, O., & Schuster, D. (2015). Discovery of novel, non-acidic mPGES-1 Inhibitors by virtual Screening with multistep protocol. *Bioorg. Med. Chem.*, *23*, 4839-4845.

Noha, S. M., Jazzar, B., Kuehnl, S., Rollinger, J. M., Stuppner, H., Schaible, A. M., . . . Schuster, D. (2012). Pharmacophore-based discovery of a novel cytosolic phospholipase A2α inhibitor. *Bioorg. Med. Chem. Lett., 22*(2), 1202-1207. doi: https://doi.org/10.1016/j.bmcl.2011.11.093

Pandey, A. V., Kempná, P., Hofer, G., Mullis, P. E., & Flück, C. E. (2007). Modulation of human CYP19A1 activity by mutant NADPH P450 oxidoreductase. *Mol. Endocrinol., 21*(10), 2579-2595. doi: 10.1210/me.2007-0245

Praxmarer, L. (2011). *Structure- and ligand-based pharmacophore modeling for mineralocorticoid receptor ligands.* (Mag. pharm.), Leopold-Franzens Universität Innsbruck, Innsbruck. Retrieved from https://permalink.obvsg.at/AC08501098

Rhöse, S. G. (2010). *Pharmacophore modeling of antiproliferative compounds: estrogen receptor subtype-selective ligands and steroid 5α reductase inhibitors.* (Mag. pharm.), Leopold-Franzens Universität Innsbruck, Innsbruck. Retrieved from https://bibsearch.uibk.ac.at/AC08352758

Rollinger, J. M., Schuster, D., Danzl, B., Schwaiger, S., Markt, P., Schmidtke, M., . . . Stuppner, H. (2009). In silico target fishing for rationalized ligand discovery exemplified on constituents of *Ruta graveolens*. *Planta Med., 75*(03), 195-204. doi: 10.1055/s-0028-1088397

Samandari, E., Kempná, P., Nuoffer, J.-M., Hofer, G., E. Mullis, P., & E. Flück, C. (2007). Human adrenal corticocarcinoma NCI-H295R cells produce more androgens than NCI-H295A cells and differ in 3β-hydroxysteroid dehydrogenase type 2 and 17,20 lyase activities. *J. Endocrinol., 195*(3), 459-472. doi: 10.1677/JOE-07-0166

Schaible, A. M., Filosa, R., Temml, V., Krauth, V., Matteis, M., Peduto, A., . . . Werz, O. (2014). Elucidation of the molecular mechanism and the efficacy in vivo of a novel 1,4-benzoquinone that inhibits 5-lipoxygenase. *Br. J. Pharmacol., 171*(9), 2399-2412. doi: 10.1111/bph.12592

Schuster, D., Kowalik, D., Kirchmair, J., Laggner, C., Markt, P., Aebischer-Gumy, C., . . . Adamski, J. (2011a). Identification of chemically diverse, novel Inhibitors of 17 beta hydroxysteroid dehydrogenase type 3 and 5 pharmacophore-based virtual screening. *J. Steroid Biochem. Mol. Biol., 125*(1-2), 148-161.

Schuster, D., Laggner, C., Steindl, T., Palusczak, A., Hartmann, R., & Langer, T. (2006). Pharmacophore modeling and in silico Screening for new P450 19 (Aromatase) inhibitors. *J. Chem. Inf. Model., 46*(3), 1301-1311.

Schuster, D., Markt, P., Grienke, U., Mihaly-Bison, J., Binder, M., Noha, S. M., . . . Wolber, G. (2011b). Pharmacophore-based discovery of FXR agonists. Part I: Model development and experimental validation. *Bioorg. Med. Chem., 19*(23), 7168-7180.

Schuster, D., Waltenberger, B., Kirchmair, J., Distinto, S., Markt, P., Stuppner, H., . . . Wolber, G. (2010). Predicting cyclooxygenase inhibition by three-dimensional pharmacophore profiling. Part I: Model generation, Validation and applicability in ethnopharmacology. *Mol. Inf., 29*(1-2), 75-86. doi: 10.1002/minf.200900071.

Temml, V., Garscha, U., Romp, E., Schubert, G., Gerstmeier, J., Kutil, Z., . . . Schuster, D. (2017). Discovery of the first dual inhibitor of the 5-lipoxygenase-activating protein and soluble epoxide hydrolase using pharmacophore-based virtual screening. *Sci. Rep., 7*, 42751.

Temml, V., Kaserer, T., Kutil, Z., Landa, P., Vanek, T., & Schuster, D. (2014). Pharmacophore modeling for COX-1 and -2 Inhibitors with LigandScout in comparison to Discovery Studio. *Future Med. Chem., 6*(17), 1869-1881.

Udhane, S. S., Parween, S., Kagawa, N., & Pandey, A. V. (2017). Altered CYP19A1 and CYP3A4 activities due to mutations A115V, T142A, Q153R and P284L in the human P450 oxidoreductase. *Front. Pharmacol., 8*(580). doi: 10.3389/fphar.2017.00580

von Grafenstein, S., Mihaly-Bison, J., Wolber, G., Bochkov, V. N., Liedl, K. R., & Schuster, D. (2012). Identification of novel liver X receptor activatos by structure-based modeling. *J. Chem. Inf. Model, 52*(5), 1391-1400. doi: 10.1021/ci300096c

Vuorinen, A., Engeli, R. T., Leugger, S., Bachmann, F., Akram, M., Atanasov, A. G., . . . Schuster, D. (2017). Potential antiosteoporotic natural product lead compounds that inhibit 17β-hydroxysteroid dehydrogenase type 2. *J. Nat. Prod., 80*(4), 965-974. doi: 10.1021/acs.jnatprod.6b00950

Vuorinen, A., G., N. L., Odermatt, A., M., R. J., & Schuster, D. (2013). Pharmacophore model refinement for 11β-hydroxysteroid dehydrogenase inhibitors: search for modulators of intracellular glucocorticoid concentrations. *Mol. Inf., 33*(1), 15-25. doi: 10.1002/minf.201300063

Waltenberger, B., Garscha, U., Temml, V., Liers, J., Werz, O., Schuster, D., & Stuppner, H. (2016). Discovery of potent soluble Epoxide hydrolase (sEH) Inhibitors by pharmacophore-based virtual screening. *J. Chem. Inf. Model., 56*, 747-762. doi: 10.1021/acs.jcim.5b00592

Walzl, M. (2010). *Molecular modeling-based analysis of 5-lipoxygenase inhibitors : common feature pharmacophore modeling and virtual screening.* (Mag. pharm.), Leopold-Franzens Universität Innsbruck, Innsbruck. Retrieved from https://permalink.obvsg.at/AC08316831

Wetzel, M., Marchais-Oberwinkler, S., Perspicace, E., Möller, G., Adamski, J., & Hartmann, R. W. (2011). Introduction of an electron withdrawing group on the hydroxyphenylnaphthol scaffold improves the potency of 17β-hydroxysteroid dehydrogenase type 2 (17β-HSD2) inhibitors. *J. Med. Chem., 54*(21), 7547-7557. doi: 10.1021/jm2008453
